# A Sensory Pathway that Controls Bacterial Zorbing

**DOI:** 10.64898/2026.07.01.735801

**Authors:** J Gosai, A Arasteh, EC Henderson, Z Yang, R Bhola, N Zeytuni, S Choubey, A Shrivastava

## Abstract

How cells transition between distinct modes of collective organization is a pivotal question in biology. We uncover the regulatory landscape governing the choice between bacterial swarming and zorbing. In *Flavobacterium johnsoniae*, zorbs are spherical, biofilm-like microcolonies that migrate collectively, whereas swarms are multilayered and ordered populations that expand planarly. We identify RgzAB, an unorthodox sensor kinase–response regulator pair governing this transition; suppressor and knockout mutants establish the importance of RgzA phosphorelay and RgzB as a downstream component. Transcriptomic and perturbation analyses link this switch to iron availability, connecting environmental sensing to collective behavior. Remarkably, RgzA activity also governs spatial organization within mixed populations: activated cells encapsulate wildtype neighbors into segregated co-zorbs, within which wildtype cells assemble into networks undergoing an abundance-dependent, percolation-like transition. Together, these findings reveal that one sensory circuit governs bacterial collectives at nested scales, from motility choice to the internal architecture of the collective.

## INTRODUCTION

Most bacteria, despite being unicellular, routinely engage in collective behaviors, of which collective motility is one of the most consequential. On soft agar surfaces, bacterial swarms are typically driven by flagella; however, individual cells do not move directly on the agar but instead swim within a thin liquid film covering the surface. Within this film, neighboring cells align and coordinate their motion, giving rise to large-scale swarms^1–3^. On relatively hard agar surfaces, where flagellar swarming is restricted, type IV pili-driven twitching motility and gliding motility can similarly generate coordinated collective movement. In these systems, cells align with one another and organize into multicellular rafts, vortices, or larger-scale swarms that migrate collectively across the surface^4–6^. While a single organism can exhibit more than one such mode, the behaviors described above run on separate, dedicated machineries. For example, in *Pseudomonas aeruginosa* flagella power swarming on soft agar while type IV pili drive collective motion on harder agar surfaces^5,7^. In such cases, the choice between behaviors can be resolved by which apparatus the cell deploys and potentially the stiffness of the surface which the cell needs to navigate. Notably, most of these coordinated collective motilities propel the bacterial population in an orderly, alignment-based arrangement of individual cells^5,8^.

*Flavobacterium johnsoniae*, a model gliding bacterium from the phylum Bacteroidota, provides a paradigm-shifting example in which the same motility machinery drives two fundamentally distinct forms of collective behavior: swarming on agar^9,10^, and zorbing on stiffer surfaces^11^. *F. johnsoniae* employs the Type IX Secretion System (T9SS) to perform gliding and has been long known to utilize this gliding apparatus for planar expansion as a quasi-2D swarm^9,10^. Recently, the same T9SS and gliding machinery was found to drive a second, strikingly different collective mode called zorbing where individual cells coalesce into three-dimensional, biofilm-like microcolonies that move as cohesive units over solid surface, and merge with one another before finally dispersing completely, placing them at an intriguing intersection of motile collectives and biofilms^11^. The zorbs, unlike swarms, show no definite symmetry or global alignment between the cells and move using several cells extending downward from the microcolony and acting as “legs”. As swarming and zorbing employ the same gliding apparatus, the choice between the collective motilities must instead rely on cellular regulation. Yet how such a choice is regulated, and how a cell commits to one collective mode over another when both rely on the same machinery, has remained unknown.

Here we present RgzAB, an unorthodox sensor kinase–response regulator circuit that inversely regulates zorbing and swarming in *F. johnsoniae*. We find that RgzAB alters motility dynamics and steers population from free-living to a cohesive state promoting zorb formation and suppressing swarming. This RgzAB-mediated shift is accompanied by major transcriptomic changes around iron acquisition and utilization, while iron availability itself alters zorbing linking environmental cue to the lifestyle switch. Finally, we find that in mixed populations, RgzAB activity drives spatial segregation within co-zorbs as RgzA-active cells encapsulate WT cells. Together, these findings establish RgzAB as a sensory switch governing collective-state selection, multicellular assembly, and emergent spatial organization in a motile bacterial community.

## RESULTS

### RgzA drives the emergence of collectively motile zorbs

During routine experiments, we serendipitously identified a spontaneous mutant of the gliding bacterium *F. johnsoniae* exhibiting a colony morphology distinct from the wildtype (WT) strain (**Fig. S1**). Whole-genome sequencing revealed a singular S158R substitution in *Fjoh_3925*, hereafter termed RgzA* (*Regulator of Zorbing*), with the asterisk denoting the S158R mutation. Homology searches, protein structure modeling, and domain predictions indicated that RgzA is an unorthodox sensor kinase containing GAF (IPR003018), Histidine kinase (IPR005467) including the DHp (Dimerization and histidine phosphotransfer; IPR003661) as well as the CA (Catalytic; IPR003594), Response regulator receiver (IPR001789), and Histidine phosphotransfer domains (IPR008207) (**Fig. 1A, Fig. S2**).

**Fig. 1.**
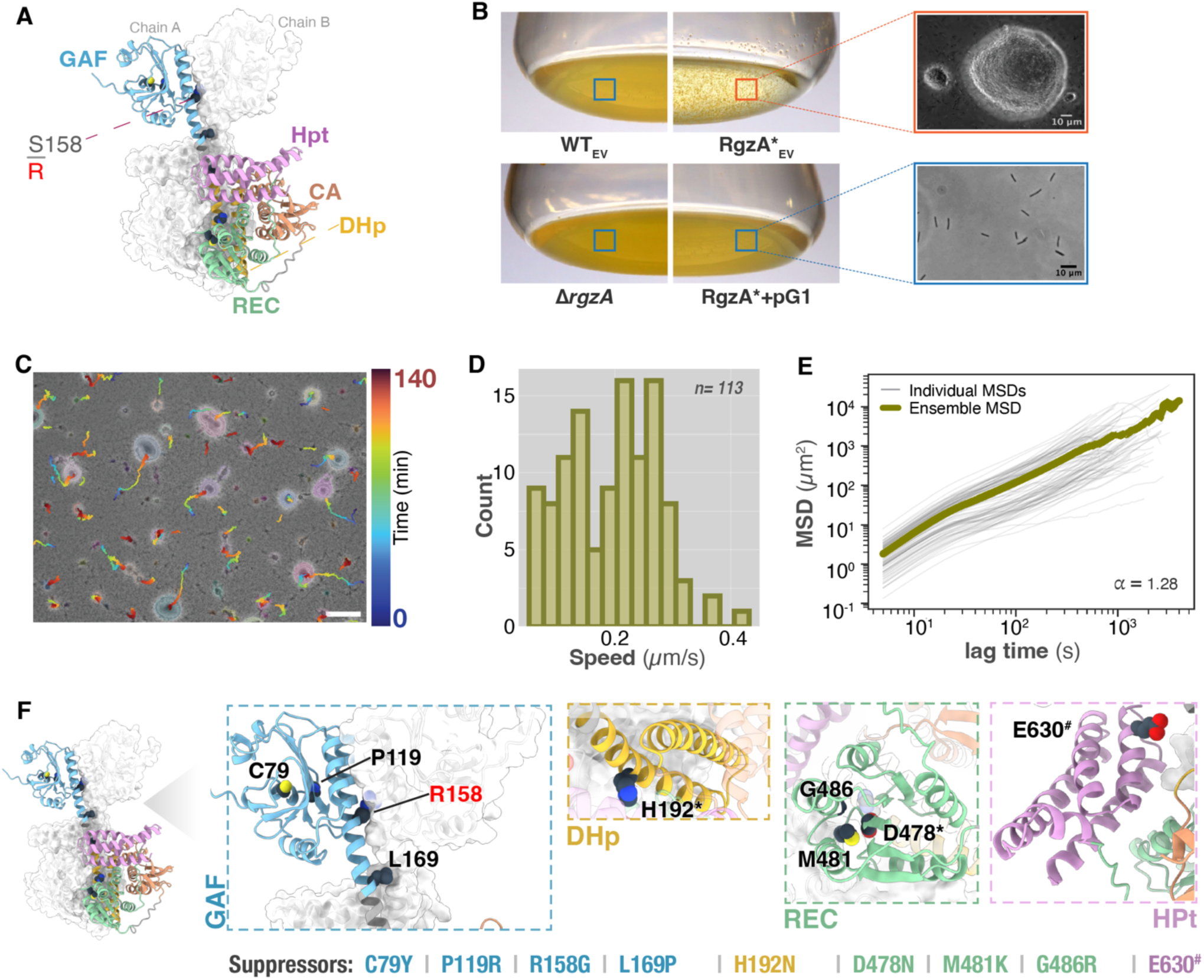
RgzA regulates formation of motile bacterial zorbs. **(A)** A predicted AlphaFold homodimer of RgzA* (R158) is shown with one monomer as a color-coded ribbon and the other as a translucent surface. On one of the chains, the predicted domains – GAF, DHp (dimerization and histidine phosphotransfer), CA (catalytic), REC (response regulator receiver), and Hpt (histidine phosphotransfer) are annotated. Ser158 in the wild type is substituted by Arg in RgzA* (S158R; shown in red). **(B)** Images of the bottoms of 125 ml Erlenmeyer flasks show the formation of large, visible zorbs by RgzA* cells. In contrast, homogeneous planktonic growth is observed in WT cells, in the *ΔrgzA* strain, and in RgzA* complemented with a wildtype copy of rgzA (pG1). Insets show representative phase-contrast images of indicated strains. Subscript EV indicates strains containing the Empty Vector-pCP23. **(C)** Time-colored trajectories of motile RgzA* zorbs overlaid on the final frame of a 140 min section of a time-lapse movie (**Movie S1**). Segmented zorbs are outlined with a colored halo. Scale bar, 100 µm. **(D)** Distribution of zorb speeds (n = 113). The median [interquartile range] is 0.20 [0.13–0.25] µm/s. **(E)** Mean squared displacement (MSD) plotted as a function of lag time (τ), showing individual trajectories (light gray) and the ensemble average (eMSD, dark olive). The anomalous diffusion exponent (α) was obtained from linear regression of log(eMSD) versus log(τ) to be 1.28 and is consistent with active motion. **(F)** Residues for which suppressor mutants were isolated are shown as spheres and annotated with their WT identity in the respective insets. Mutations in the GAF domain (blue) localize to the predicted ligand-binding pocket or the dimerization interface, including the original RgzA* mutation (R158). Active site residues denoted by asterisks (*), including H192 in the DHp domain (yellow) and D478 in the REC domain (green). Truncation of the HPt domain (light magenta) via a stop codon at E630 (denoted by #) also results in suppression (Specific mutations are listed below the dimer and described in **Table S4**).

Interestingly, under the standard growth conditions in culture flasks (see Methods), RgzA* cells grew as large, visible-to-naked-eyes aggregates ranging from tens- to hundreds of microns in diameter. Complementation with a WT copy in RgzA* background reverted to WT-like phenotype. We created the deletion mutant, Δ*rgzA*, which, surprisingly resembled the WT and stayed planktonic as well (**Fig. 1B, S3).** This indicated that RgzA* was an activating mutation. Dimer modeling using Alphafold3 confidently predicted that RgzA forms a homodimer (**Fig. 1A, S4**). The model showed that in the RgzA* variant, the Arg158 sidechain extends close to Asp30 of the opposing monomer, unlike Ser158 in WT RgzA (**Fig. S5**). This contact provides electrostatic stabilization of the homodimer (**Fig. S5**). Phenotypic findings along with structural modeling indicate that RgzA* is an activating mutation.

Visualizing in real time in a simple flow-cell on a phase contrast microscope, we found that the RgzA* aggregates formed, merged and moved as collectives on the glass surface (**Movie S1, Fig. 1C**). Segmentation and tracking revealed that the RgzA* collectives moved with a mean squared displacement exponent of ∼1.3, consistent with active motility and similar to that previously described for zorbs formed by WT *F. johnsoniae* (**Fig. 1D, E**). Together, these features identify the RgzA* collectives as zorbs. Notably, whereas WT *F. johnsoniae* zorbs were originally observed under low-nutrient conditions, in an underoil microfluidic device^11^, RgzA* cells formed zorbs abundantly independent of nutrient concentrations and under simple flow-cell conditions **(Fig. S6).** This observation, combined with the constitutive zorb phenotype of RgzA* and the WT-like behavior of the *ΔrgzA* strain, confirms that RgzA governs zorbing and that the RgzA* variant induces the zorb-forming state. Notably, this constitutively zorbing strain makes the study of zorbing highly accessible by providing a genetic handle to dissect the regulation of this unique form of collective behavior.

To gain more insight in the working of RgzA, we then performed suppressor mutagenesis in the RgzA* strain (see **Fig. S7** for details) and isolated suppressors that revert to WT-like swarming phenotype. The mutations spanned all of the domains of RgzA and all reverted to the WT-like phenotypes **(Fig. S7, Table S4)**. Specifically, mutations were identified at predicted ligand-binding or proximal residues within the GAF domain (C79Y, P119R; **Fig. 1F**; also see **Fig. 4A**) and at residues associated with predicted dimerization interfaces (R158G, L169P). Additional suppressor mutations were identified in the conserved active site residues of the histidine kinase domain (H192D) and in the response regulator active site (D478M) (**Fig. S8**) that are known to be crucial for phospho-relay in other histidine kinases and response regulators^12–14^. Mutations M481K and G486R also mapped in the proximity of D478, and a truncation mutation (E630#), removing the final ∼30 amino acids of the histidine phosphotransfer domain, was likewise recovered (**Fig. 1C inset**). This indicated that RgzA* mediated activation of zorbing required action of each of the domain including the dimerization interfaces and internal phosphorelay residues.

Additionally, taxonomic survey for similar proteins indicated that RgzA homologs are broadly distributed within the original genus and the phylum Bacteroidota with relatively sparse presence in Pseudomonadota, Cyanobacteriota and other phyla. **(Fig. S9).** Together, the genetic, phenotypic, and structural analyses suggest a model where ligand binding, dimerization, and internal phosphorelay precede RgzA activation, and that the S158R substitution may stabilize the dimer, locking the protein in an active state. Consistent with this model, only the RgzA* variant elicits zorbing and associated phenotypes, whereas deletion of *rgzA* is indistinguishable from WT.

### RgzA drives zorbing through coupled adhesion and motility control

Consistent with the intercellular coherence seen in the zorbs, and with their biofilm-like nature, we found that RgzA* exhibited early and amplified biofilm formation at both solid–liquid interfaces (walls of polystyrene 96-well plates) and solid–liquid–air interfaces (glass tube rims) as compared to the WT **(Fig. 2A, B).** It is likely that the increased intercellular cohesion that possibly stabilizes zorbs also enhanced adhesion to abiotic surfaces.

**Fig. 2.**
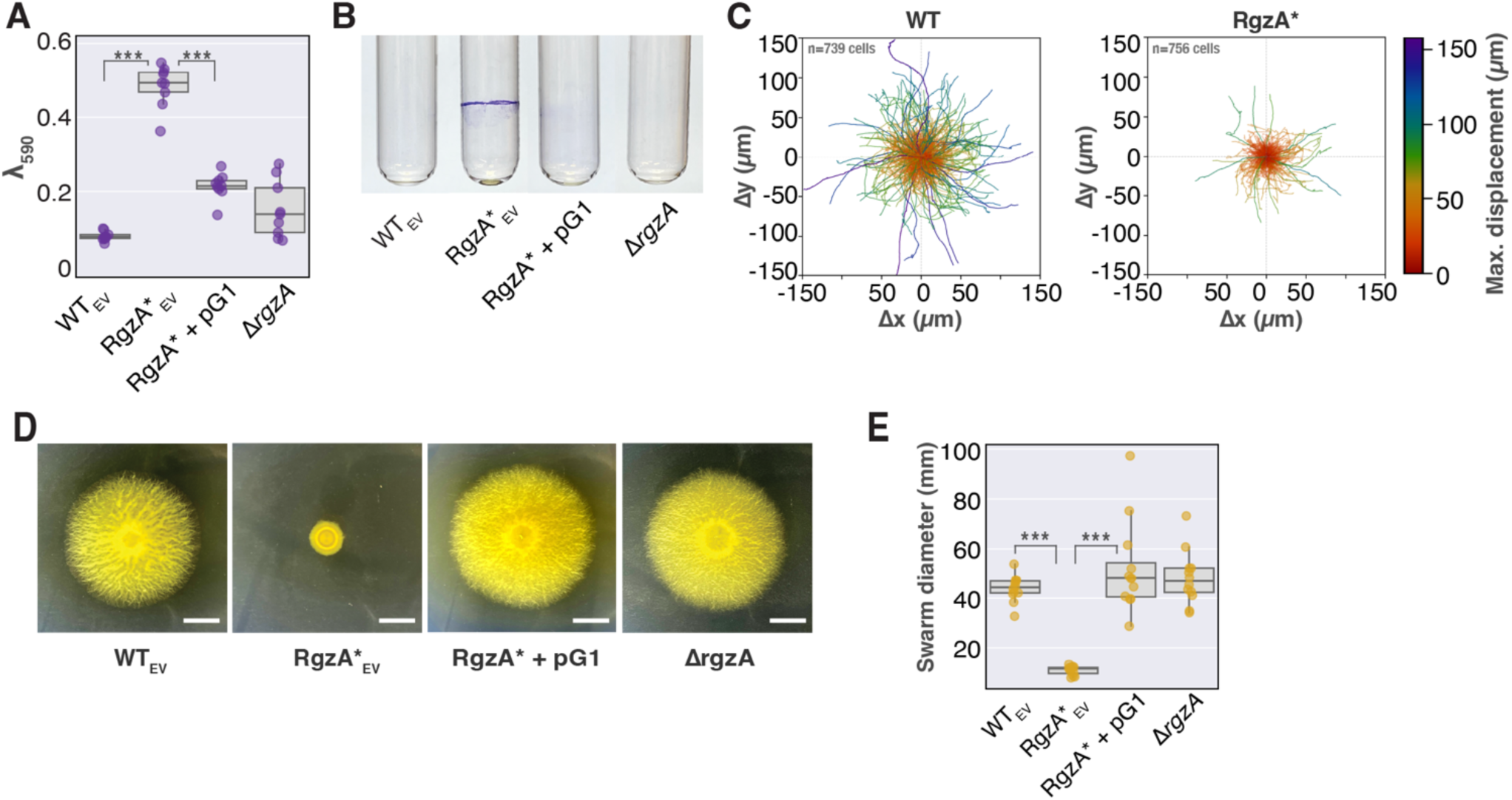
RgzA increases biofilm formation, alters single cell motility and suppresses swarming. (**A)** Quantification of biofilm formation on polystyrene plates. The plot shows crystal violet absorbance at 590 nm following biofilm staining and ethanol solubilization (n = 9 per strain; three biological repeats with three technical repeats each). (**B)** Photographs show crystal violet staining on glass culture tubes after 24 h of growth. A biofilm ring is visible at the air-liquid-solid interface for RgzA*, while no biofilm formation is observed in other tested strains. (**C)** Single cell trajectories of WT and RgzA* represented as rose plots. Individual cell paths are normalized to a common origin and color-mapped to maximum displacement (distance to the farthest point in the trajectory). WT cells navigate longer distances, whereas RgzA* movement is more localized. n = 739 (WT) and 756 (RgzA*). (Also see **Movie S2** and **Fig. S10**). (**D)** RgzA* results in diminished swarming motility. While the WT swarms to form an expanded colony on agar, RgzA* fails to swarm. Exogenous expression of WT-copy of *rgzA* (complementation) restores the WT phenotype, while the *rgzA* deletion mutant shows no difference from WT. Scale bar = 10 mm. (**E)** Quantification of swarm diameters shows a significant difference between WT and RgzA*. N = 12 per strain (three biological repeats with four technical repeats each). In panels A and E, the datapoints on the plot show individual observations, box shows median line flanked by interquartile ranges (IQR), whiskers show 1.5x of the IQR. *** denotes *P <* 0.001 by randomized block ANOVA followed by Tukey’s post-hoc test. p-value comparisons between all groups are available in **Table S7**. Subscript EV indicates strains containing the Empty Vector pCP23.

In many bacteria, a shift toward sessility in terms of amplified biofilm formation also results in reduced motility^15–18^. In addition to zorbing, *F. johnsoniae* can traverse solid surfaces as individual cells as well as in a swarm. We tested if RgzA* led to changes in either single-cell or swarming motility. Microscopic time-lapse imaging and tracking of individual motility, however, revealed that RgzA* cells retained individual motility but showed considerable difference in the patterns as compared to the WT. RgzA* cells predominantly exhibited oscillatory motion, reversing more frequently than WT cells, leading to reduced overall displacements (**Fig. 2C, S10, S11 and Movie S2, S3)**. The differences in the motility are corroborated by a significant shift in the rotational bias of the motor associated with the Type IX secretion system, which drives gliding in *F. johnsoniae*. Prior tethering assays of sheared WT cells showed that ∼92% of motors rotate counterclockwise (CCW)^19,20^, whereas in RgzA* cells only ∼67% of T9SS motors rotated CCW (**Fig. S12**).

Remarkably, unlike individual gliding motility, RgzA* cells exhibited substantially reduced collective swarming on agar. RgzA* shows a significant reduction in radial expansion as compared to the wild type (**Fig. 2D, E**). In contrast, both the complemented strain and Δ*rgzA* displayed swarm expansion comparable to WT. This suppression of swarming may reflect a shift in how cells organize collectively: rather than dispersing across the surface as multilayered, aligned, and oriented swarms, RgzA* cells preferentially organize into three-dimensional, cohesive zorbs. Together, these data indicate that RgzA activation enhances adhesion and alters motility dynamics, driving a population-level shift toward zorb formation while suppressing swarming. Thus, RgzA appears to function as a regulatory switch that biases the same gliding population toward distinct modes of collective organization and motility.

### RgzA forms a signaling circuit with an adjacent response regulator

Adjacent to *rgzA* (*Fjoh_3925*), lies the gene *Fjoh_3924 (rgzB,* hereafter), predicted to encode a single domain response regulator (RR) (**Fig. 3A**). We hypothesized, based on the genomic neighborhood, that they could make a functional pair. Structural modeling of dimeric RgzA in complex with an RgzB monomer using AlphaFold3^21^ confidently predicted an interaction at the interface between the RgzA Hpt domain and RgzB (**Fig. 3B, S4E**). The model further indicated that the predicted active site residues *i.e.*, His607 in RgzA-Hpt and Asp51 in RgzB are positioned in close proximity (**Fig. 3B**). In order to test if RgzB is required for RgzA-mediated phenotypes, we deleted *rgzB* in the RgzA* background. We found that this strain abolished RgzA*-associated phenotypes of zorbing and biofilm formation thus restoring this strain to WT-like behavior. In contrast, deletion of *rgzB* in WT background showed no difference from the WT phenotypes. Conversely, complementing the strain lacking RgzB in RgzA* background with a WT copy of *rgzB* led to restoration to parent RgzA*-like phenotypes including robust zorbing, amplified biofilm formation, and diminished swarming (**Fig. 3C-F**). These results indicate that RgzA and RgzB form a functional gene circuit and that RgzB acts downstream of RgzA.

**Fig. 3.**
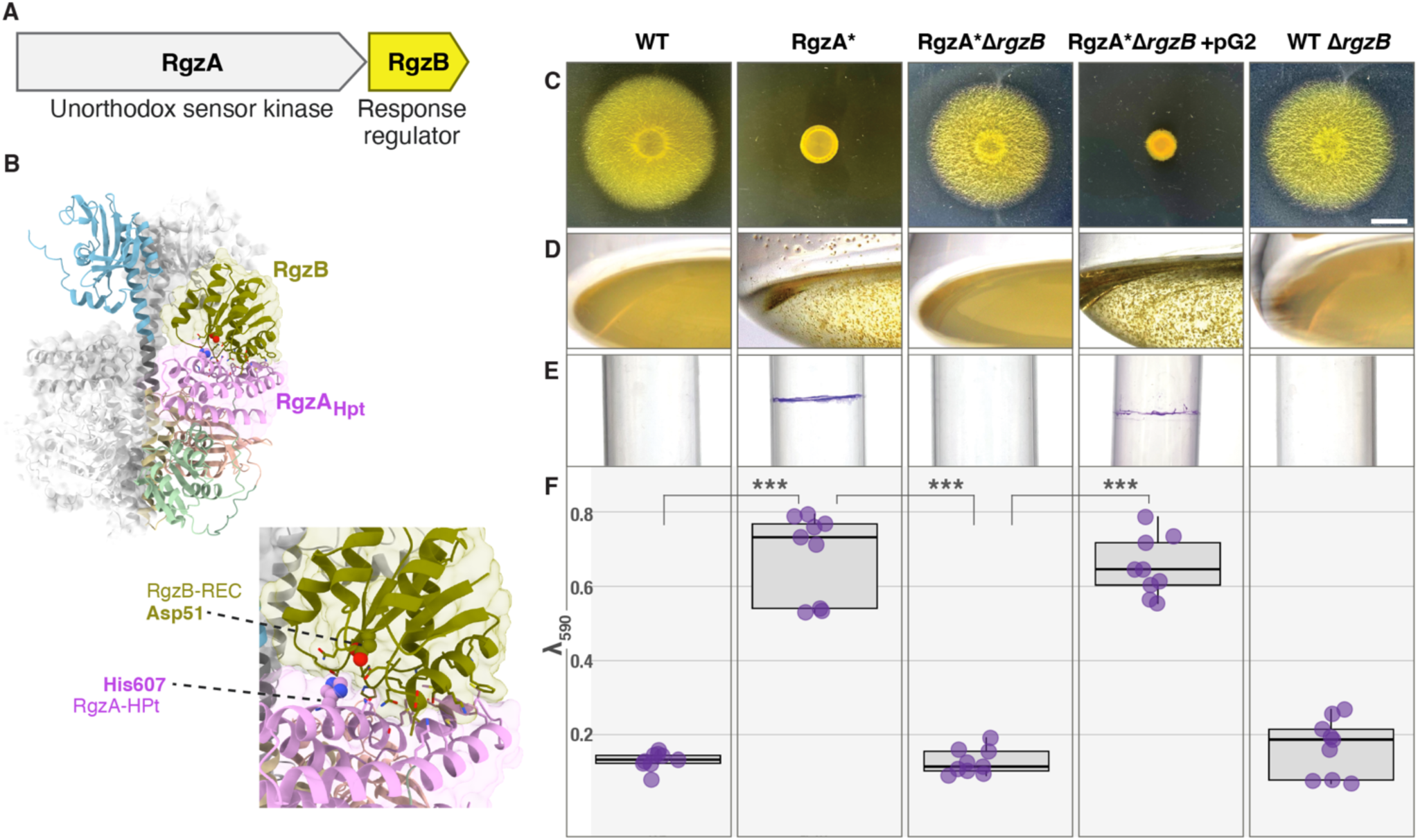
RgzB is functionally linked to RgzA and controls zorbing, swarming, and biofilm formation. (**A)** An illustration indicating that the genes encoding RgzA (*Fjoh_3925*) and RgzB (*Fjoh_3924*) are next to one another on the genome. (**B)** Predicted complex of RgzA and RgzB. An AlphaFold3 model of an RgzA homodimer (gray and color-coded by domain) in complex with an RgzB (colored olive) is shown. The model places RgzB in close proximity to the RgzA HPt domain. Residues within 3Å of the RgzA-HPt/RgzB interface are shown via a stick representation. The predicted donating His and receiving Asp residues of RgzA-HPt and RgzB, respectively, are highlighted as spheres. Lower panel: A zoomed-in view illustrates the proximity between the predicted phosphotransfer residues. **(C–F)** Phenotypic characterization of WT, RgzA*, and *rgzB* mutant strains. Assays include swarming motility **(C)**, zorb formation **(D)**, biofilm formation on glass **(E)**, and biofilm formation in polystyrene plates **(F)**. Deletion of *rgzB* in the RgzA* background (RgzA Δ*rgzB*) results in the loss of all RgzA*-associated phenotypes, restoring a WT-like state. Complementation with a plasmid-borne copy of *rgzB* (from pG2) in this background restores the RgzA* phenotypes. Deletion of *rgzB* in the WT background (WT Δ*rgzB*) yields no detectable phenotypic differences from WT. Scale bar in panel C = 10 mm. In panel F, the datapoints on the plot show individual observations n = 9 including three biological and three technical repeats, box shows median line flanked by interquartile ranges (IQR), whiskers show 1.5x of the IQR. *** *P <* 0.001 by randomized block ANOVA followed by Tukey’s post-hoc test. p-value comparisons between all groups are available in **Table S7**.

### Iron suppresses RgzA-driven zorbing and dominates its transcriptional program

Structural analysis of the predicted RgzA model revealed an oval-shaped pocket centrally located within the GAF domain, suggestive of a potential ligand-binding site (**Fig. 4A**). The binding pocket features a bottleneck approximately 2.8 Å wide and an internal cavity measuring ∼4.4 Å in diameter (**Fig. 4B, C**), dimensions consistent with coordination of divalent cations. Metal ion-binding predictions performed using the Metal Ion-Binding site prediction and docking server^22^ (MIB2) identified copper, iron and zinc as the most likely candidate cationic ligands for this site (**Table S5**). Five common metal-binding residues, His50, Ser52, Cys79, Tyr115 and Cys132 are situated next to the core of the binding pocket in close proximity and are predicted to participate in metal coordination (**Fig. 4D**).

**Fig. 4.**
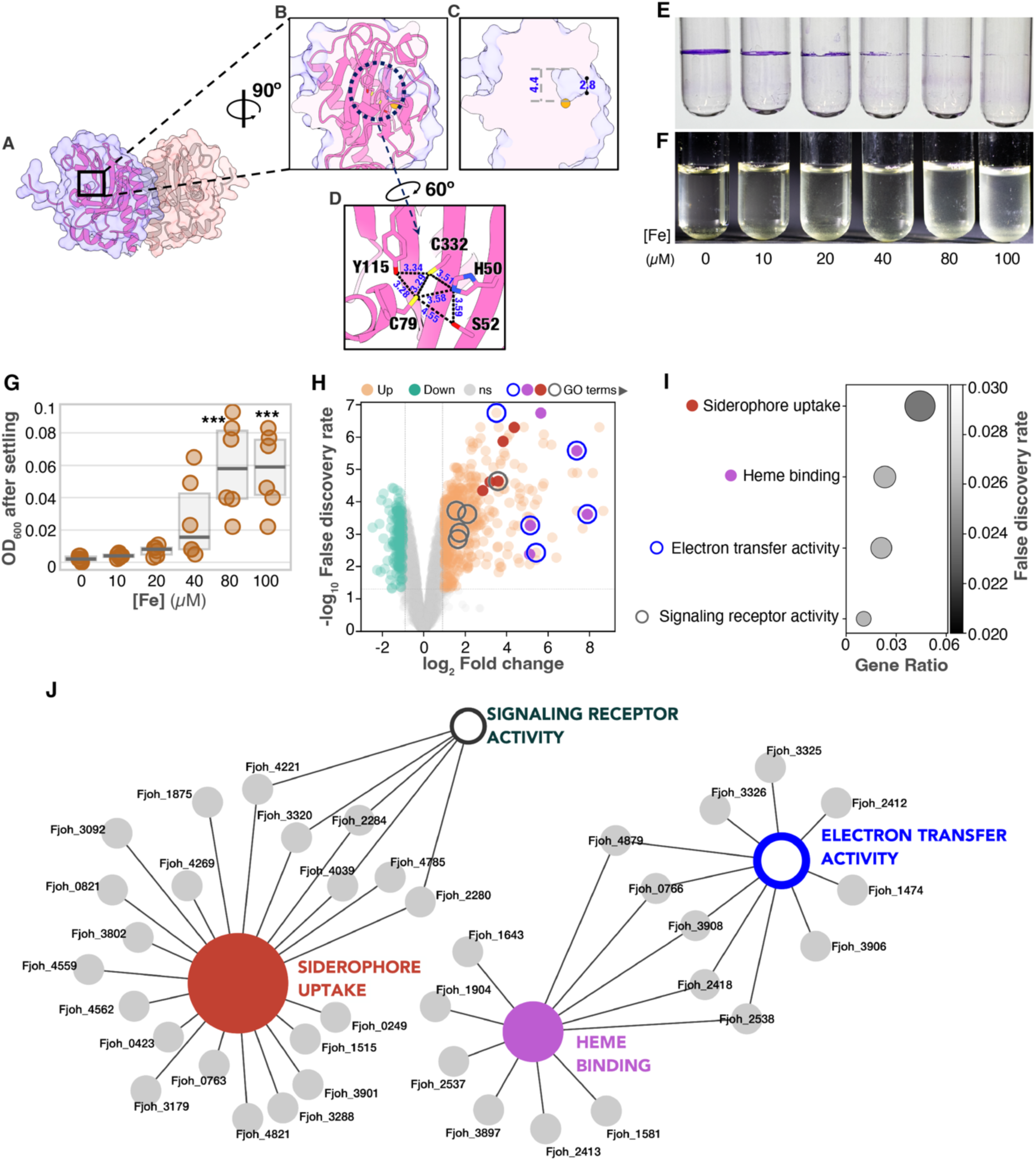
RgzA-mediated zorbing is iron-linked. (**A)** Predicted ligand-binding pocket is highlighted with a black square on the predicted dimeric structure of RgzA GAF domains. (**B)** Enlarged side view of the oval-shaped binding pocket is shown depicting sectioned molecular surface density (purple) superimposed on the cartoon representation (pink) of the GAF domain. The predicted divalent cation is shown in orange, and key residues implicated in ligand binding are encircled with black dotted lines. **(C)** Side view of the binding pocket. The molecular surface (purple) is sectioned to reveal the internal dimensions of the pocket, with the divalent cation shown in orange. **(D)** Close-up view of residues His50, Ser52, Cys79, Tyr115, and Cys132 positioned in proximity to the cation. Hydrogen bonds are represented by black dashed lines. Blue colored numbers denote distances in angstrom. **(E)** Representative photographs of glass culture tubes demonstrating an iron-dependent reduction in biofilm formation at the air-liquid-solid interface. **(F)** Increasing iron concentrations reduce zorb formation. Zorbs were grown in Erlenmeyer flasks and transferred to tubes for the ease of photography. Strong zorbing leads to biomass sedimenting to the bottom, leaving the upper region less turbid while reduced zorbing results in uniformly high turbidity throughout the tube. **(G)** Quantification of turbidity in the upper region of the culture tubes shown in (F), measured as OD₆₀₀. The datapoints on plot show 6 individual observations, box shows median line flanked by interquartile ranges (IQR), whiskers show 1.5x of the IQR (n = 6 per condition); *** *P* < 0.001 by randomized block ANOVA followed by Dunett’s post-hoc test. **(H)** RNA-seq analysis identified a large set of differentially expressed genes. The volcano plot shows 483 significantly upregulated genes (orange) and 169 significantly downregulated genes (teal), with 3,953 non-significant genes (grey). Additional annotations on the figure include enriched GO terms which are described in subsequent panels. Dashed lines indicate thresholds of | log₂FC | > 1 and FDR < 0.05. **(I)** Dot plot showing significantly enriched Gene Ontology (GO) terms among differentially expressed genes (DEGs). Dot size represents the gene ratio (proportion of DEGs annotated to each term), and grayscale intensity corresponds to the false discovery rate. **(J)** Gene-concept network (cnetplot) illustrating the association between enriched GO terms and their constituent DEGs. Enlarged colored nodes represent GO terms; smaller grey nodes represent individual genes labelled with their respective locus tags. Edges connect genes to their associated GO terms.

To test the in silico predictions, and whether metal availability influences zorbing, we assessed the effect of divalent cations fitting the dimension criteria on zorbing. Separately, CuCl_2_, NiCl_2_, ZnCl_2_, and FeCl_3_ as well as CaCl2 and MgCl_2_ were added to cultures during zorb formation. Among these, only FeCl_3_ significantly reduced zorb formation in RgzA* cells, whereas the other metals had no detectable effect (**Fig. S13**). Iron addition also markedly reduced biofilm formation (**Fig. 4E, S14**) and zorb formation (**Fig. 4F, G**) across a range of iron concentrations.

In order to resolve the scope of regulation downstream of RgzA, we performed RNA-seq analysis on RgzA* zorbs and similarly grown WT cells. The analysis revealed differential expression of approximately 700 genes, representing ∼15% of the total genes in *F. johnsoniae*, indicating that RgzA activation is associated with broad metabolic and lifestyle shifts accompanying zorb formation (**Fig. 4H, S15, Table S6**). Gene set enrichment analysis of these differentially expressed genes showed significant enrichment of iron-related Gene Ontology (GO) terms, including strong upregulation of iron acquisition and respiration-associated pathways. Notably, multiple genes encoding siderophore systems and iron–sulfur cluster– associated functions were upregulated to a remarkably high degree (**Fig. 4 H-J**). These transcriptomic findings are consistent with the biofilm formation and zorbing response to iron addition, suggesting iron availability might modulate RgzA activity and that iron abundance likely shifts RgzA toward a less active state leading to inhibition of zorbing.

### The RgzA activity state dictates spatial organization in co-zorbs

Having established that RgzA activity determines whether *F. johnsoniae* adopts a dispersed swarming state or assembles into cohesive zorbs, we next asked whether the RgzA signaling state also determines how cells organize *within* the collective. Magesh and coworkers previously showed that zorbing WT *F. johnsoniae* can encapsulate other bacterial species to form multispecies co-zorbs^23^. To further investigate how RgzA influences collective organization within a mixed microbial population, we co-cultured WT and RgzA* cells. Remarkably, the two populations spontaneously segregated into spatially distinct compartments within the same co-zorb, with WT cells preferentially occupying the core and RgzA* cells forming an encapsulating outer layer (**Fig. 5A, B**). This is also consistent with the observation that in larger co-zorbs, likely resulting from the fusion of smaller co-zorbs, multiple WT cores could be seen within a shared RgzA* shell **(Fig. 5C**). Interestingly, this spatial segregation was present at a range of mixing ratios, including when the relative proportion of RgzA* was dropped to one-tenth of the WT (**Fig. S16**). These findings indicate that encapsulation may be an intrinsic outcome of zorbing and that RgzA-driven zorbing can drive spatial segregation even within an otherwise isogenic population.

**Fig. 5.**
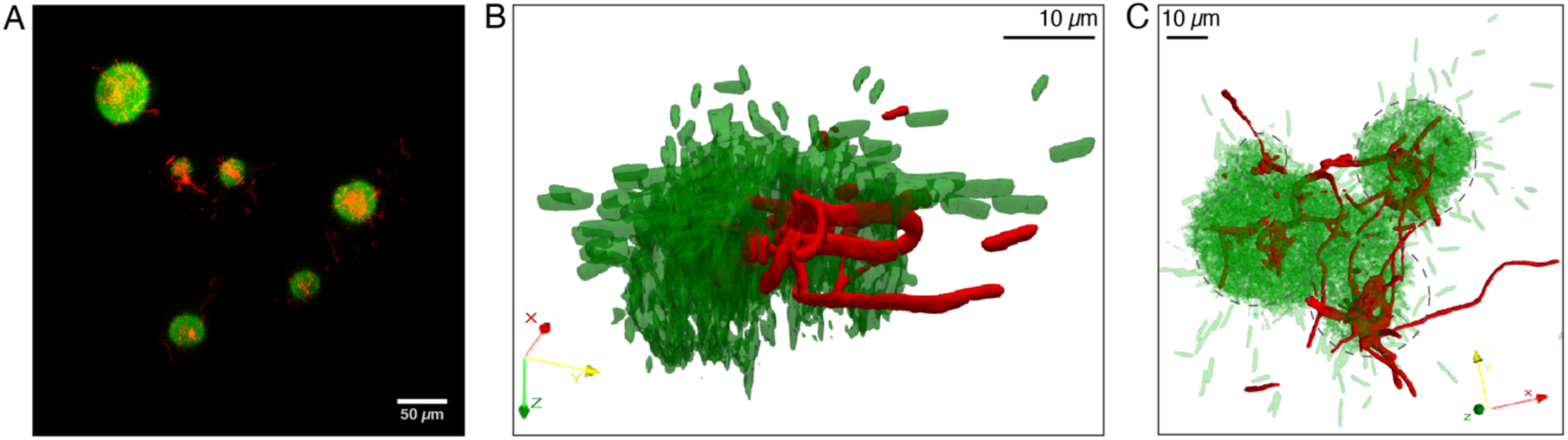
RgzA* cells encapsulate WT cells in a co-culture. **(A)** Micrograph of mixed co-zorbs of WT (red) and RgzA* (green). WT largely occupies the insides of the co-zorb while also extending elongated cellular extensions outward of the encapsulating RgzA*. **(B)** A 3D render of a representative co-zorb. A quarter of the green RgzA* volume has been sliced off to show the core of the co-zorb which shows volume occupied by the red (WT) biomass. Cellular extensions can also be seen to be extending outside the boundaries of RgzA* cell volume. **(C)** A large co-zorb likely formed from merging of smaller co-zorbs shows that each of the latter ones have their own core of WT with cells.

Confocal imaging of RgzA*–WT co-zorbs at higher resolution revealed that the encapsulated WT cells did not remain as an amorphous bundle but instead formed tangled networks with extended cellular projections, which appeared to connect cellular clusters into lattice-like structures spanning large regions of the co-zorb interior (**Fig. 5C, S16**). This network-like organization was consistently observed regardless of zorb size or WT:RgzA* mixing ratio, suggesting that some degree of internal connectivity is an intrinsic, emergent feature of the encapsulated state. This spatial organization notably differs from that observed in earlier multispecies co-zorbs, in which WT *F. johnsoniae* preferentially localized to the periphery. Thus, activation of RgzA not only promotes zorb formation but also reshapes the spatial organization of cells within a mixed population, thus raising the question of what, in turn, shapes the specific architecture of the encapsulated network itself.

### Within RgzA*-encapsulated co-zorbs, WT abundance tunes internal network architecture

To test how relative population sizes influence the architecture of the network-forming encapsulated WT subpopulation, we co-cultured WT and RgzA* strains across mixing ratios ranging from WT:RgzA* = 1:2 to 10:1. As mentioned earlier, across all conditions, the clustering was observed (**Fig. 6A–C, S15**). While WT cells were consistently encapsulated at every mixing ratio, the internal organization of the WT biomass varied strongly with relative population abundance.

**Fig. 6.**
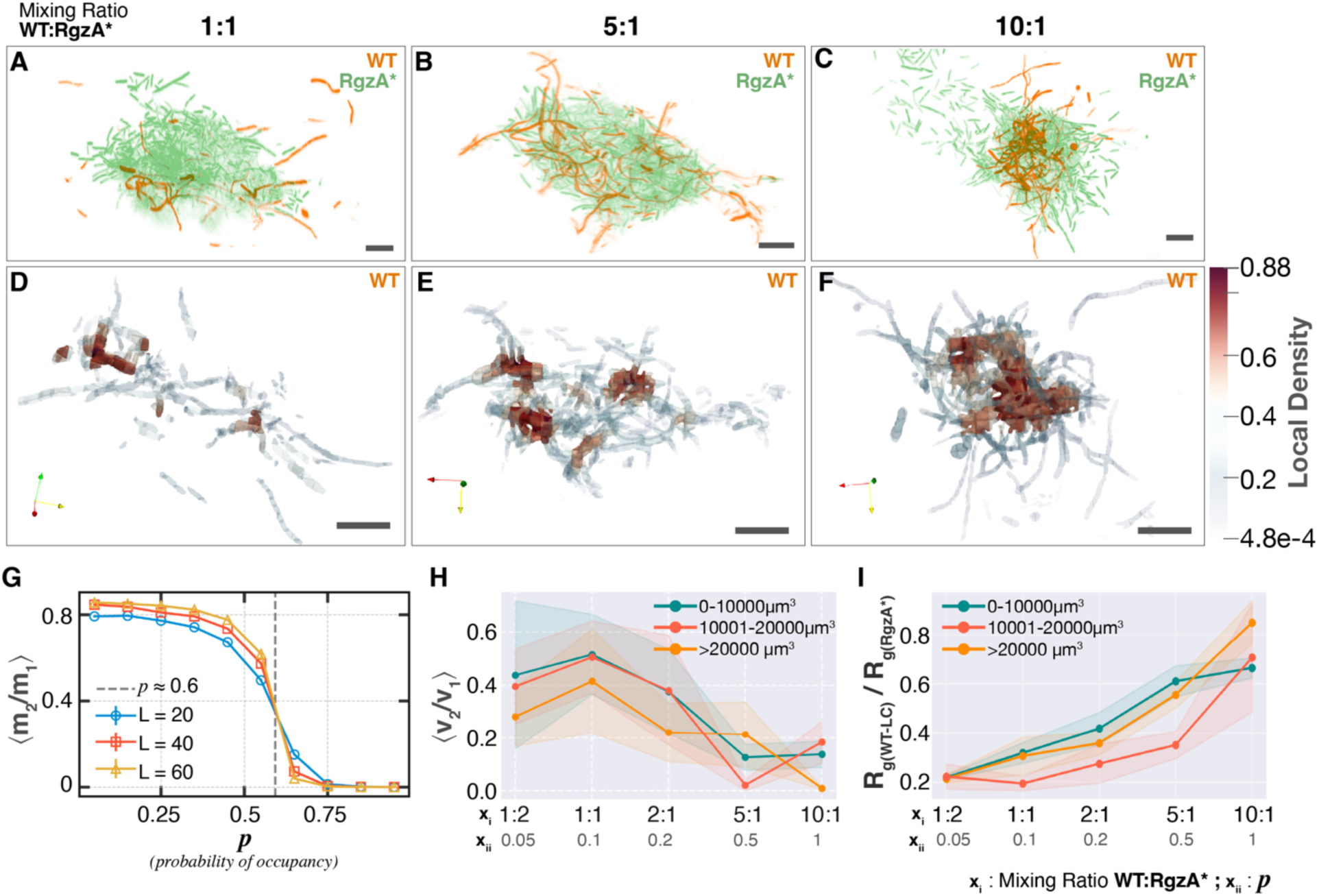
WT cells within RgzA*-WT co-zorbs undergo a percolation-like transition. **(A-C)** Representative 3D perspective views of confocal micrographs showing WT (orange) and RgzA* (green) co-zorbs at the indicated mixing ratios (WT:RgzA*). Scale 10 µm. **(D-F)** Three-dimensional renderings of the WT biomass encapsulated within the RgzA* co-zorb shown in panels A-C color-mapped by local density. The dense regions serve as nodes for percolation analysis. Here, the RgzA* channel is not shown to allow clarity in visualization of WT networks. At higher mixing ratios, WT cells consolidate from dispersed, low-density extensions (cool tones) into high-density clusters (warm tones). Scale 10 µm. **(G)** Percolation transition simulated on a 2D square lattice. The ratio of the second-largest cluster mass to the largest cluster mass (⟨m_2_/m_1_⟩) is plotted against site occupancy probability. The dashed vertical gray line marks the theoretical percolation threshold (*p_c_* ≈ 0.6). Colors represent lattice size, where the number of sites is (L^2^), with (L) ranging from 20 to 60. **(H)** The volume ratio of the second largest to the largest WT cluster (⟨v_2_/v_1_⟩) from WT-RgzA* co-zorbs is plotted as a function of mixing ratio (x_i_) and the derived occupancy probability (x_ii_). Shaded regions represent standard error. The ratio across different zorb size classes (colored lines) recapitulates the characteristic pattern seen in simulated percolation (as seen in panel G). **(I)** Spatial extent of the largest WT cluster. The normalized radius of gyration, defined as the radius of gyration of the largest WT cluster (R_g(WT-LC)_) divided by R_g_ of the RgzA* (R_g(RgzA*)_) biomass is plotted as a function of mixing ratios and occupancy probability (*p*) for the same volume classes as panel G. With increasing *p*, the largest cluster occupies a progressively greater fraction of the total available volume which is indicative of percolation-driven cluster growth. For panels **H and I**: n = 24, 20, and 35 for classes of zorb sizes <10000, 10001-20000, and >20000 µm^3^ respectively.

At low relative proportions, WT biomass organized into multiple small clusters resembling a nodal network with extended cellular projections. As relative WT abundance increased, these clusters began to consolidate, eventually approaching a single dominant cluster (**Fig. 6D–F**), independent of overall co-zorb size (**Fig. 6H, I**). This abundance-dependent consolidation resembles the classic signature of a percolation transition^24^, a framework from statistical physics describing how connectivity-based aggregation shifts as the density of a component increases thus providing us with a quantitative ‘lens’ to analyze the transition observed here.

To quantify this feature, we relied on the ratio of the volumes of the second-largest and largest WT cluster^25–27^ (〈v₂/v₁〉) within a network. For a network undergoing a percolation transition, this ratio declines in an inverted sigmoidal fashion as individual clusters progressively consolidate. Consistent with this, 〈v₂/v₁〉 for WT declined progressively as relative WT abundance increased (**Fig. 6H**): at low WT fractions, clusters remained discrete and comparable in size, producing relatively high and stable 〈v₂/v₁〉 values, whereas at higher WT fractions the largest cluster expanded rapidly relative to the rest, driving a sharp drop in 〈v₂/v₁〉 consistent with the emergence of a dominant cluster. Although the current statistics do not allow identification of a sharp percolation threshold (pc), the overall pattern is consistent with this behavior.

Simulation of classical percolation (**Fig. 6G**) illustrates the three regimes this behavior would be expected to traverse, depending on the occupation probability relative to the critical threshold (*p_c_*): in the subcritical regime (*p* < *p_c_*), all clusters are finite and typically small; at criticality (*p* ≈ *p_c_*), the system exhibits large fluctuations with clusters spanning a broad range of sizes and no single dominant component; and in the supercritical regime (*p* > *p_c_*), a unique macroscopic (system-spanning) cluster emerges while the remaining clusters stay finite and comparatively small.

To further test this feature, we examined radius of gyration dynamics. As with the transition described above, the radius of gyration of the largest WT cluster (normalized to overall zorb size) increased with relative WT abundance, eventually approaching values consistent with the largest cluster spanning most of the zorb interior (**Fig. 6I**). In our system, mixing ratio plays a role qualitatively similar to the occupation probability p in lattice percolation, where increasing the fraction of one component tunes connectivity and aggregation propensity. Consistent with this, the cluster-size distribution P(s) progressively shifted toward larger clusters as the mixing ratio increased **(Fig. S17 and Supplementary Text**), supporting enhanced cluster growth and connectivity. While we are constrained by finite system size and sampling, the reproducible enrichment of larger clusters across conditions supports the emergence of composition-dependent connected networks.

Together, these results reveal that a single sensory circuit choreographs collective life at three nested scales at once: RgzA activity first sets the population’s basic mode of movement (to swarm or to zorb), then, draws the boundary between a zorb’s inside and outside while the interior attains and operates in distinct ecology of its own.

## DISCUSSION

Zorbs deviate from the prevalent mutual exclusivity between biofilm formation and swarming among bacteria^11^. Interestingly, for the collective translocation, zorbs share the gliding machinery with swarming which is a completely different kind of collective motion exhibited by *F. johnsoniae*^11,28^. This demands that cellular regulation must resolve which, if any, collective motion the cell performs. Identification of the RgzAB gene circuit provides a mechanistic foundation for this switch, demonstrating how the signaling pathway directs a population away from single-cell gliding or orderly swarming and into a cohesive, motile zorb state.

Structural modeling of the multidomain sensor RgzA confidently predicted that it forms a homodimer and indicated that the S158R substitution in RgzA* stabilizes the dimer. Consistent with this, several independent observations support that RgzA* is an active state of the protein: suppressor mutations in key residues across the RgzA* protein abolish the associated phenotypes, *ΔrgzA* phenocopies the WT, and RgzA* uncouples zorbing from the nutritional conditions that constrain it in the WT. This makes RgzA* an excellent strain to easily study zorbing.

The predicted sensory domain of RgzA, GAF, is one of the most versatile sensory domains in bacterial signaling proteins, coordinating ligands that span a wide range of chemistry and physical sizes^29–31^. Modeling of the RgzA GAF domain indicated, both, a pocket of appropriate dimension, and an arrangement of residues consistent with metal-cation coordination^31–34^. In the perturbation experiments, among the cations we tested, only iron affected zorbing, and it did so by suppressing zorb formation.

Together these observations suggest a model in which the apo-(ligand-free) state of RgzA exists as an active dimer which is stabilized in RgzA* mimicking low-iron conditions and promoting zorbing. This is corroborated by low nutrient conditions required for WT *F. johnsoniae* zorbing^11^ and promotion of aggregation in low nutrient medium^35^. Binding of iron to RgzA GAF domain may then shift the equilibrium to monomeric, inactive RgzA, leading to suppression of zorbing. While most sensor kinases attain their active, dimer form upon ligand binding, our model places RgzA as an exception. It is noteworthy that such an exception is not unprecedented. A recent case that deviates from this paradigm and bears striking resemblance to the proposed RgzA model is the multidomain GAF-containing sensor kinase PdtaS in *Mycobacterium tuberculosis*. PdtaS is an active dimer in its ligand-free state, and is directly inhibited by cation binding, which disrupts dimerization^36^. Some other examples include FixL in *Ensifer meliloti*^37^ and WalK in *Staphylococcus aureus*^38^, all sensors containing GAF/PAS domains.

The internal phosphorelay of RgzA, evidenced by the loss of zorbing when key catalytic residues are mutated, culminates in RgzB activation, likely through phosphotransfer via the Hpt domain. Consistent with RgzB acting downstream of RgzA, deleting rgzB abolishes zorbing in the RgzA* background but has no effect in the WT. The phosphorelay and the circuit architecture is consistent with previously reported hybrid HK-RR pairs. RgzB is a CheY-type single domain response regulator (RR) lacking any DNA-binding output domain. Regulators of this family usually propagate the signal by interacting with a target protein directly^39^. Further studies identifying its interacting partners can shed more light on the sequence of events immediately following RgzB activation.

Activation of RgzA alters the expression of ∼15% of the ‘gene-ome’, with a strong enrichment for iron acquisition and respiratory pathways such as siderophore uptake, and electron transfer. This is also accompanied by strong upregulation of genes encoding fumarate reductase, nitrite reductase, c-type cytochromes, NiFe-hydrogenase, and a high-affinity terminal oxidase **(Table S6)**. The expression profile resembles an iron-starvation response^40–42^ and is consistent with adaptation to the oxygen-limited interior of dense yet non-motile aggregates^43–45^. In this context, the co-induction of siderophore uptake makes sense: respiration under these conditions is iron-intensive, and the cells appear to be preparing for life inside their own collective structure. The transcriptome is thus consistent with iron-linked regulation of zorbing.

Activated RgzAB circuit, ultimately suppresses swarming and promotes zorb formation. Swarming colony expansion in *F. johnsoniae* is well characterized. On nutrient-poor agar, cells glide outward in a thin film in which they are organized into aligned clusters that move co-directionally, guided by paths of secreted extracellular material laid down by leading cells^46,47^. This is a coordinated, contact-dependent 2D migration of a dense population, sharing the core physical features of swarming in flagellated bacteria^48^ but being powered by the T9SS gliding motor rather than flagella^10^. On the other hand, in zorbing, cells self-assemble asymmetrically into discrete, spherical three-dimensional microcolonies built around a polysaccharide core which traverse surface as a unit using the same T9SS/gliding machinery^11^. In this manner, the RgzAB circuit does not only switch the population from planktonic to a cohesive collective state but also determines the selection between two completely opposite modes of collective motility using the same molecular machinery.

As RgzAB sets the collective state of the population, it also determines the cell’s place within the collective. Encapsulation of WT by RgzA* cells when they are co-cultured reverses previously observed encapsulation hierarchy where WT encapsulated unrelated bacterial species^23^. This indicates that an actively zorbing population (RgzA* here) acts as the constitutive ‘*encapsulator*.’ This phenotype likely reflects the importance of coordinated changes in motility, cohesion, and matrix production. Inside these spatially segregated co-zorbs, the story becomes increasingly remarkable starting with the appearance of emergent network like form in the encapsulated WT population. The elongated, interconnected projections formed by the WT cells extending through the RgzA* shell (**Fig. 5C**) resemble in appearance the filamentous architectures seen in biofilms and other non-motile aggregates, where they are associated with altered cell–cell adhesion^49^, division defects under oxygen limitation^50^, or both. This finding fuels future investigations about how this sensory circuit may shape the spatial organization of polymicrobial communities and microbiota.

The network-like projections when visualized in co-zorbs with a varying WT:RgzA representation, were found to undergo a percolation-like transition^24^. Percolation transition describes the dynamics of independently growing clusters and them coalescing into one continuous, system-spanning network. Such connectivity can enable long-range coordination and transport that isolated clusters cannot support^51,52^. Percolation-like behavior has rarely been described in bacterial systems. Previous studies have reported such connectivity transitions as being associated with membrane-potential propagation and quorum-sensing within biofilms^53,54^. Observing such a transition in a spatially segregated, confined population is therefore notable and may reflect on the biofilm-like nature of the zorbs.

We note that the observed networks are confined within the finite dimensions of the RgzA*-derived co-zorbs, imposing a finite-size limit on cluster growth (**Fig.6H, I**). Additionally, the assemblies are captured at heterogeneous stages of development, introducing considerable variability; for example, a snapshot of newly fused co-zorbs might show multiple similarly sized clusters even at high abundance. Time-resolved visualization and tracking of emergent WT networks can aid in precisely defining the underlying transition. We observe increased cell death within the zorb core (**Fig. S18**), in line with stress-associated mortality reported in dense bacterial aggregates^55–57^ suggesting that this spatial organization may create gradients in resources and stress that differentially impact cell survival within the zorb. Future studies could resolve if such network-like organization could support metabolite exchange between otherwise disconnected regions, provide routes for coordinated dispersal, or generate spatial heterogeneity in survival.

Finally, identification of RgzA through an activating allele rather than a deletion illustrates a general limitation of screens in sensory transduction that search for a lack of phenotype: sensor kinases remain inactive until their activating signal arrives, and a deletion under standard laboratory conditions often reports only that the kinase was already inactive. Constitutively active alleles, whether engineered, as in the systematic approach of Claverie and colleagues in *Streptococcus*^58^ or serendipitous, as in this work, decouple the regulon from the unknown upstream signal and expose circuits that deletion approaches miss.

While this work identifies RgzAB as a regulator of collective motility in gliding Bacteroidota and establishes a mechanistic, biophysical, and ecological framework for zorbing, it also provides a foundation for exploring the broader biology and ecological significance of this unusual collective behavior. Future studies can now investigate the range of environmental and physiological signals sensed by RgzA and map its spatiotemporal activation as cellular segregation emerges within the colony. The RgzAB system also offers an opportunity to connect sensory signaling directly to the mechanics of collective motility by determining how RgzA activation coordinates changes in T9SS motor dynamics and how these changes culminate in zorb formation. Likewise, defining how signals are transmitted downstream of RgzB, which lacks an obvious output domain, should reveal additional components of this regulatory pathway. The encapsulated cellular networks provide another exciting avenue for investigation, particularly for determining whether their spatial organization promotes signaling, dispersal, survival, or other adaptive functions under conditions that more closely resemble *Flavobacterium*’s natural environments, such as those explored by Magesh and colleagues^59^.

Importantly, RgzA* now provides a robust and experimentally tractable system for studying the biology and physics of bacterial collective organization and motility. Together, our findings establish RgzAB as a sensory circuit that governs transitions among single-cell gliding, swarming, and zorbing and, in doing so, determines how microbial populations organize themselves in space. This framework creates new opportunities to uncover how sensory regulation, motility, and cell–cell interactions are integrated to generate distinct collective states and, ultimately, how these states enable microbial populations to navigate, adapt, and compete in complex environments.

## MATERIALS AND METHODS

### Bacterial strains and culture conditions

Bacterial strains, plasmids and primers used in this study are listed in supplementary **Tables S1-S3**. *Flavobacterium johnsoniae* CJ1827 (WT) and its derivatives were grown in Casitone Yeast Extract (CYE) medium (10 g/L casitone, 5 g/L yeast extract, 10 mM Tris-HCl pH 7.5; 1.5% agar for solid medium) at 30°C. For motility assays, *F. johnsoniae* Motility Medium (FJMM) (one-third strength CYE broth) was used. Swarming assays were performed on PYE2 agar (2 g/L peptone, 0.5 g/L yeast extract, 1.0% agar)^60^. *Escherichia coli* strains were cultured in Luria-Bertani (LB) medium at 37°C. When required, antibiotics were added at the following concentrations Ampicillin (100 μg/mL), Erythromycin (100 μg/mL), Tetracycline (20 μg/mL), Streptomycin (100 μg/mL).

### Assessment of single cell motility and Zorb formation

Single cell motility is assayed as described previously ^60^. Briefly, overnight culture (100 µL) of *F. johnsoniae* cells was inoculated in 5 ml of FJMM in a 125 ml Erlenmeyer flask and grown for 6-8h at 25°C shaken at 80 rpm. Cells grown in motility culture conditions as described above are washed in FJMM employing gentle centrifugation at 3000g for 3 min. Washed cells were then loaded into a tunnel slide (double-sided tape spacers between a slide and coverslip), incubated for 5 min, and flushed with 200 µL of FJMM. Live imaging was performed using a Nikon Eclipse 50i phase contrast (Nikon Corporation, Japan) microscope equipped with CMOS camera (CS165MU1, Thorlabs, USA) at 15 frames per second (fps). Resultant movies were analyzed using RABiTPy^61^ to assess the motility dynamics of single cells. When RgzA* cells are cultured in Motility culture conditions as described as above, they grow almost exclusively as large visible-to-naked-eyes zorbs. To quantify the extent of zorbing, 3 mL of culture was allowed to settle in a glass culture tube for 5 min. The OD600 from gently aspirated top 200 µL was measured to determine the degree of aggregation.

### Time-lapse imaging of zorbs and computational analysis

To assay zorbing dynamics for longer time period, a flow cell (FC81, Biosurface technologies) connected to a syringe pump (Pump 22, Harvard Apparatus) was mounted on inverted phase contrast microscope Nikon Diaphot equipped with CMOS camera (DCC1545M, Thorlabs, USA). Inoculation of RgzA* cells was done via single injection at the beginning using one of the ports of the flow cell. After a 5-min incubation to allow initial adherence of cells, unadhered ones were removed by a 1-min flush at a fast flow rate (25 mL/min), followed by a constant flow of FJMM (100µL/min). Time-lapse images were captured over 12–16 h using the above-described microscope and camera. Only the frames with actively motile zorbs were used for quantitative estimation of motility dynamics. For this, image processing was performed in Fiji^62^; segmentation of zorbs was achieved using a custom pixel classification model trained in ilastik version 1.4.1rc2 ^63^; tracking was performed using TrackMate 7 with ilastik used as the detector^64^. Resultant trajectories were analyzed using a custom Python pipeline to calculate velocities and mean squared displacement (MSD).

### Genetic manipulation of *F. johnsoniae*

The general process essentially followed previously published method^65^. In brief, around 2-kb genomic fragments each upstream and downstream of the target gene were PCR-amplified using Phusion polymerase (New England Biolabs, MA) and cloned into *BamHI/SalI*-digested pRR51 via Gibson Assembly (New England Biolabs, MA). The resulting suicide vector was introduced into *F. johnsoniae* via biparental conjugation with *E. coli* S17-1 λ*pir*. Allelic exchange was performed using a two-step recombination strategy: initial selection on erythromycin followed by counter-selection on streptomycin to isolate second-crossover events. PCR-amplification and DNA sequencing were used to confirm the genotypes. For expression of a gene exogenously, region spanning the target gene was amplified using PCR and cloned in KpnI/BamHI digested pCP23 using directional restriction cloning. This construct is then transferred to *F. johnsoniae* using *E. coli* S17-1 λ*pir* and transconjugants are selected on tetracycline.

### Recreation of the RgzA* (S158R) mutation

During the deletion of *porX* (*Fjoh_2906*) in *F. johnsoniae*, a spontaneous mutant displaying a non-spreading, convex, and shiny colony morphology was isolated. Whole-genome sequencing (SeqCenter, Pittsburgh, PA) identified a point mutation in *Fjoh_3925* (AGC → AGA), resulting in an S158R substitution. This strain was designated FJASU_27. To recreate this mutation in a wildtype background, a 2-kb genomic fragment encompassing the mutation site was PCR-amplified from FJASU_27 using primers PASU_286 and PASU_287. This fragment was cloned in pRR51 followed by processes as described above. The resulting point mutant in the WT background was confirmed using long-read sequencing (Plasmidsaurus, USA), designated as FJASU_28 and was designated as the RgzA* strain.

### Suppressor mutagenesis

Five microliters of overnight grown culture of RgzA* was spotted on a PYE2 plate for swarming assay as described above (without the dilution). Simply letting the incubation past 48 h allowed for spontaneous occurrences of suppressor mutants presenting as flares coming out as mini-swarm arcs from circumference of the colony. Biomass from the edge of these flares was carefully picked up, cultured and used for isolation of genomic DNA followed by PCR amplification and subsequent long-read sequencing of *Fjoh_3925* region (Plasmidsaurus, KY, USA).

### Swarming plate assays

Qualitative and quantitative analysis of swarming of *Flavobacterium* strains was performed as follows. An overnight culture of the respective strain in CYE broth was prepared starting from an isolated colony on a plate. The resultant culture was diluted to OD_600_ of 0.1. Five microliters of this diluted suspension of bacteria were spotted in the center of a PYE-2 plate (see Bacterial strains and culture conditions). The plates were incubated at 25°C for 48 h in a 100% relative humidity chamber and were photographed then with an Apple iPhone. The diameters of the swarms were measured using Fiji^62^ and plotted. Several measurements of diameter on each of the swarm image were measured and averaged. At least three technical as well as three biological repeats were performed.

### Biofilm formation

Quantitative biofilm formation was studied for *F. johnsoniae* strains on 96-well polystyrene plates as per the method described in ^66^. Briefly, overnight cultures were inoculated in 96 well plates at OD_600_ of 0.05 and incubated for 24 h. At the end of incubation, the plate was gently shaken, and the culture was transferred to a fresh plate to obtain planktonic OD600. The original plate was washed three times with water to remove unadhered cells, and air-dried. The biofilm was then stained with 0.1% crystal violet (CV) for 45 minutes. Bound CV was dissolved using 95% Ethanol solution and was read calorimetrically on a plate reader (Spectramax M5, Molecular Devices, USA).

Photographic and qualitative estimation of biofilm on air-liquid-solid interface was done in glass culture tubes. Overnight culture of *F. johnsoniae* strains was used to inoculate 3 ml CYE medium in a 15 ml culture tube. These were shaken at 180 rpm at 30°C for 24 h. At the end of incubation, the culture was discarded, and unbound cells were washed with two successive plain water rinses; washed tubes were dried and stained with 0.1% CV and dried again before being photographed using smartphone camera against bright white background to capture the presence or thickness of the biofilm ring.

### Estimation of direction of motor rotation

The rotational bias of the T9SS gliding motor was determined as previously described using a shearing and tethering assay, and rotational speed and direction were analyzed using custom scripts described previously^19,20^.

### Structural modeling

Prediction of the dimeric and monomeric 3D structures of RgzA and in complex with RgzB were performed using Alphafold2^67^ and domain predictions by InterPro^68^ SMART prediction tool^69^. Structural models of dimeric RgzA were generated using the AlphaFold2 pipeline with the multimer model preset to predict protein complexes for the latter. The full-length amino acid sequence of the protein (UniProt ID: A5FCZ0) was used as input. Predictions were performed locally on Compute Canada using the full database configuration and the multimer model. The following sequence and structure databases were used for template search and multiple sequence alignment (MSA) construction: UniRef90, UniRef30, BFD, MGnify, PDB70, and PDB mmCIF structures. The top-ranked model, based on predicted local distance difference test (pLDDT) and predicted aligned error (PAE) metrics, was selected for structural interpretation. All structural visualizations, domain mapping, and interface analyses were performed using UCSF ChimeraX2^70^. To identify putative metal-binding sites, the AlphaFold2-predicted models were analyzed using the Metal Ion-Binding Site Prediction and Docking Server (MIB2)^22^. Output from MIB2 was used to infer the identity of likely coordinating residues and candidate divalent cations (e.g., Fe²⁺, Zn²⁺, Cu2⁺). Structural visualization and annotation were performed using UCSF ChimeraX2.

### Assessment of effect of various cations on RgzA* phenotypes

Based on the prediction of size of ligand binding pocket (See **Fig. 4**), cations were tested to see if they affected the extent of zorbing. WT or RgzA* cells were cultured in CYE overnight. This culture was used to start motility culture conditions – which included none or 100 µM of either of FeCl_3_, ZnCl_2_, NiCl_2_, CuCl_2_, MgCl_2_ or CaCl_2_. At the end of incubation, zorbing was photographed and assayed quantitatively as described above. Based on these results, a range of concentrations of FeCl_3_ (10-100 µM) was tested for its effect on zorb and biofilm formation as described above.

### RNAseq analysis

*F. johnsoniae* WT and RgzA* were grown in Motility conditions (see culture conditions above) and the biomass from each of them was harvested. RNA-seq was performed on RgzA* vs WT; n = 3 biological replicates of each strain. Total RNA was isolated using RNeasy kit from Qiagen, USA. RNA was sent to SeqCenter (SeqCenter, Pittsburgh, PA) for RNA-seq analysis. Briefly, the process was as follows. The isolated RNA was subjected to ribodepletion. Quality control and adapter trimming was performed with bcl-convert (a proprietary Illumina software). Read mapping was performed with HISAT2^71^. Read quantification was performed using Subread’s featureCounts^72^ functionality. Read counts loaded into R and were normalized using edgeR’s Trimmed Mean of M values (TMM) algorithm^73^. Subsequent values were then converted to counts per million (cpm). Reads were aligned to genome of *Flavobacterium johnsoniae* UW101 from NCBI accession number NC_009441. Differential expression analysis was performed using edgeR’s Quasi-Linear F-Test (qlfTest) functionality against treatment groups. Significantly differentially expressed (DE) genes were determined with the thresholds —|logFC| > 1 and FDR < .05. The differentially expressed gene’s normalized counts per million were then used to create a heatmap (**See Fig. S15**). Volcano plot was plotted using this data using custom python script employing matplotlib.

### Gene set enrichment analysis

Gene Ontology (GO) Molecular Function enrichment analysis was performed using a hypergeometric test with Benjamini-Hochberg multiple testing correction, implemented in custom Python script using the SciPy ^74^ and pandas^75^. GO term annotations were assigned using PANNZER^76^. Differentially expressed genes were tested for enrichment against a background of all detected genes. Terms with a Benjamini-Hochberg adjusted p-value (FDR) < 0.05 were considered significantly enriched. Results were visualized as a dot plot in which the x-axis represents the GeneRatio (number of differentially expressed genes annotated to a term divided by the total number of differentially expressed genes), dot size reflects the number of annotated genes, and dot shading reflects the FDR value. Figures were generated in Python using matplotlib. To visualize gene-to-term relationships, a cnetplot network was constructed for the 4 significantly enriched GO Molecular Function terms. Edges were defined between genes and their annotated GO terms, yielding a bipartite graph of 41 nodes and 47 edges. Node and edge tables were exported in Python and imported into Cytoscape (v3.10.1) for visualization and layout.

### Confocal laser scanning fluorescence microscopy of RgzA* and mixed culture zorbs

Monoculture RgzA* zorbs were grown as described above (*Motility culture conditions*) using RgzA* carrying pAS43-GFP. The zorb suspension was transferred to a 30 mm polystyrene petri dish and allowed to settle for 3–5 minutes, after which the supernatant was decanted. Attached zorbs were gently resuspended in saline (0.85% sodium chloride w/v) and mounted in tunnel slides for imaging. Imaging was performed on a Zeiss LSM 880 confocal laser scanning microscope (CLSM) or Nikon AX R Confocal microscope using either a 60x oil-immersion or 20x air objective. For mixed culture zorbs, overnight cultures of either FJASU_101 (*F. johnsoniae* expressing mStrawberry) or FJASU_81(*F. johnsoniae* expressing GFP), and non-fluorescent RgzA* or FJASU82 (*F. johnsoniae* RgzA* expressing GFP) were each harvested at OD 1.0. Cultures were mixed at ratios of WT-strain:RgzA*-strain = 1:2, 1:1, 2:1, 5:1, and 10:1, to a total volume of 100 µl, which was used to inoculate 5 mL of FJMM which was then grown as in motility conditions described above. These were harvested and imaged as described in the above paragraph.

### Segmentation and quantitative image analysis of CLSM micrographs of zorbs

The resultant z-stacks from imaging performed as above were processed in napari^77^ in combination with custom python scripts. First, the planktonic cells that were outside the zorbs were removed from the raw images which were saved as a multi-channel OME-tiff files. These were used in BiofilmQ^78^ for a segmentation and quantification. The resultant data was exported as csv and vtk file formats. vtk files for visualization were exported and used in Paraview^79^ for visualization if needed. csv files were used in the custom python pipeline to determine density thresholds and perform clustering. Clustering was done using a binary clustering method based on connected component analysis implemented in custom python pipeline. Resultant clusters were used for percolation analysis. Percolation was tested using matrics such as ratios of volumes of second largest to that of the largest cluster at a range of mixing ratios. Radius of gyration was measured from the same dataset and normalized using the radius of gyration of the encapsulating strain (RgzA*) biomass.

### Percolation simulation

The study was conducted on L × L square lattice with periodic boundary conditions, where each simulation began with a square lattice in which all sites were initially occupied. Site dilution was then performed by randomly removing sites from the lattice with probability 1-*p*, where *p* represents the occupation probability. Cluster identification was performed using a breadth-first search (BFS) based burning algorithm that systematically identifies all connected components in the diluted lattice. The algorithm begins by marking all occupied sites in the diluted lattice as unvisited, then selects an unvisited occupied site as a seed point for cluster identification. From this seed site, the algorithm explores all connected nearest neighbors using 4-connectivity in two dimensions through a queue-based breadth-first traversal. All sites reached during this BFS expansion are assigned to the same cluster and marked as visited. This process is repeated iteratively until all occupied sites have been visited and assigned to their respective clusters. The BFS algorithm ensures that each identified cluster represents a maximal connected component of occupied sites, where nearest-neighbor relationships on the square lattice define connectivity.

For each diluted lattice configuration, we tracked all identified clusters and recorded their sizes, defined as the number of sites per cluster. We specifically focused on M₁, the size of the largest cluster, and M₂, the size of the second-largest cluster. The ratio M₂/M₁ was calculated for each lattice realization to track the percolation transition^25–27^. To obtain statistically robust results, we performed ensemble averaging over multiple independent realizations. For each occupation probability p, we generated N independent random diluted lattices, typically ranging from 1000 to 2000 samples. Each sample was individually analyzed using the BFS clustering algorithm to obtain M₁ and M₂ values. The M₂/M₁ ratio was computed for each sample, and the ensemble average <M₂/M₁> was calculated across all N samples.

The occupation probability *p* was systematically varied across the parameter space to map the behavior of <M₂/M₁> as a function of site occupation density. The parameter range was carefully chosen to capture the percolation transition region, where the most significant changes in cluster structure occur and where the interplay between the largest and second-largest clusters becomes most pronounced. Lattice sizes ranged from L = 20 to L = 60 to investigate finite-size effects and ensure results were representative of the thermodynamic limit.

## ACKNOWLEDGEMENTS

AS was supported by National Institutes of Health – National Institute of General Medical Sciences award R35GM147131 and National Science Foundation MCB award 2530162. NZ was supported by Natural Sciences and Engineering Research Council of Canada Discovery Grants (RGPIN/03031-2022). NZ is a member of the Centre de Recherche en Biologie Structurale (CRBS), funded by Fonds de Recherche du Québec (Health Sector) Research Centres Grant #288558. R.B. was funded by the Deutsche Forschungsgemeinschaft (DFG, German Research Foundation) - SFB 1551, Project-ID 464588647.

## AUTHOR CONTRIBUTIONS

JG and AS conceptualized this study. JG collected and analyzed the majority of the data. AA contributed to some of the genetic manipulations and phenotypic characterizations. ECH performed the cell tethering assays and contributed to some of the genetic manipulations. ZY and NZ performed the structural modeling. RB and SC developed the phase transition and percolation simulations and contributed to the analysis of associated experimental data. JG and AS wrote the manuscript.

## DATA AVAILABILITY

Custom Python codes for image and data analysis are available on GitHub at https://github.com/jitendragosai/sensory-origins-of-zorbing. Transcriptomic data is deposited on NCBI Gene Expression Omnibus Accession GSE329926. The raw data and codes are deposited on Zenodo at DOI: 10.5281/zenodo.20033030.

## REFERENCES

1. Wu, Y., and Berg, H.C. (2012). Water reservoir maintained by cell growth fuels the spreading of a bacterial swarm. Proceedings of the National Academy of Sciences 109, 4128–4133. 10.1073/pnas.1118238109.

2. Wu, Y., Hosu, B.G., and Berg, H.C. (2011). Microbubbles reveal chiral fluid flows in bacterial swarms. Proceedings of the National Academy of Sciences 108, 4147–4151. 10.1073/pnas.1016693108.

3. Harshey, R.M. (2003). Bacterial Motility on a Surface: Many Ways to a Common Goal. Annual Review of Microbiology 57, 249–273. 10.1146/annurev.micro.57.030502.091014.

4. Kaiser, D. (1979). Social gliding is correlated with the presence of pili in Myxococcus xanthus. Proc Natl Acad Sci U S A 76, 5952–5956. 10.1073/pnas.76.11.5952.

5. Burrows, L.L. (2012). Pseudomonas aeruginosa Twitching Motility: Type IV Pili in Action. Annual Review of Microbiology 66, 493–520. 10.1146/annurev-micro-092611-150055.

6. Nakane, D., Odaka, S., Suzuki, K., and Nishizaka, T. (2021). Large-Scale Vortices with Dynamic Rotation Emerged from Monolayer Collective Motion of Gliding Flavobacteria. Journal of Bacteriology 203, 10.1128/jb.00073-21. https://doi.org/10.1128/jb.00073-21.

7. Rashid, M.H., and Kornberg, A. (2000). Inorganic polyphosphate is needed for swimming, swarming, and twitching motilities of Pseudomonas aeruginosa. Proceedings of the National Academy of Sciences 97, 4885–4890. 10.1073/pnas.060030097.

8. Kaiser, D. (2007). Bacterial Swarming: A Re-examination of Cell-Movement Patterns. Current Biology 17, R561–R570. 10.1016/j.cub.2007.04.050.

9. Chang, L.E., Pate, J.L., and Betzig, R.J. (1984). Isolation and characterization of nonspreading mutants of the gliding bacterium Cytophaga johnsonae. J Bacteriol 159, 26– 35.

10. McBride, M.J. (2001). Bacterial Gliding Motility: Multiple Mechanisms for Cell Movement over Surfaces. Annu. Rev. Microbiol. 55, 49–75. 10.1146/annurev.micro.55.1.49.

11. Li, C., Hurley, A., Hu, W., Warrick, J.W., Lozano, G.L., Ayuso, J.M., Pan, W., Handelsman, J., and Beebe, D.J. (2021). Social motility of biofilm-like microcolonies in a gliding bacterium. Nat Commun 12, 5700. 10.1038/s41467-021-25408-7.

12. Wolanin, P.M., Thomason, P.A., and Stock, J.B. (2002). Histidine protein kinases: key signal transducers outside the animal kingdom. Genome Biol 3, reviews3013.1. 10.1186/gb-2002-3-10-reviews3013.

13. Gao, R., Mack, T.R., and Stock, A.M. (2007). Bacterial Response Regulators: Versatile Regulatory Strategies from Common Domains. Trends Biochem Sci 32, 225–234. 10.1016/j.tibs.2007.03.002.

14. Jenal, U., and Galperin, M.Y. (2009). Single-domain response regulators: molecular switches with emerging roles in cell organization and dynamics. Curr Opin Microbiol 12, 152–160. 10.1016/j.mib.2009.01.010.

15. Gosai, J., Anandhan, S., Bhattacharjee, A., and Archana, G. (2020). Elucidation of quorum sensing components and their role in regulation of symbiotically important traits in *Ensifer* nodulating pigeon pea. Microbiological Research 231, 126354. 10.1016/j.micres.2019.126354.

16. Rossi, E., Paroni, M., and Landini, P. (2018). Biofilm and motility in response to environmental and host-related signals in Gram negative opportunistic pathogens. J. Appl. Microbiol. 125, 1587–1602. 10.1111/jam.14089.

17. Ruhal, R., and Kataria, R. (2021). Biofilm patterns in gram-positive and gram-negative bacteria. Microbiological Research 251, 126829. 10.1016/j.micres.2021.126829.

18. Chen, X., Wang, L., and Sourjik, V. (2025). Swimming or sessile: the interplay between c-di-GMP signalling and flagellar motility. Current Opinion in Microbiology 87, 102632. 10.1016/j.mib.2025.102632.

19. Shrivastava, A., Lele, P.P., and Berg, H.C. (2015). A Rotary Motor Drives *Flavobacterium* Gliding. Current Biology 25, 338–341. 10.1016/j.cub.2014.11.045.

20. Trivedi, A., Miratsky, J.A., Henderson, E.C., Singharoy, A., and Shrivastava, A. (2025). A molecular conveyor belt-associated protein controls the rotational direction of the bacterial type 9 secretion system. mBio 16. 10.1128/mbio.01125-25.

21. Abramson, J., Adler, J., Dunger, J., Evans, R., Green, T., Pritzel, A., Ronneberger, O., Willmore, L., Ballard, A.J., Bambrick, J., et al. (2024). Accurate structure prediction of biomolecular interactions with AlphaFold 3. Nature 630, 493–500. 10.1038/s41586-024-07487-w.

22. Lu, C.-H., Chen, C.-C., Yu, C.-S., Liu, Y.-Y., Liu, J.-J., Wei, S.-T., and Lin, Y.-F. (2022). MIB2: metal ion-binding site prediction and modeling server. Bioinformatics 38, 4428–4429. 10.1093/bioinformatics/btac534.

23. Magesh, S., Schrope, J.H., Soto, N.M., Li, C., Hurley, A.I., Huttenlocher, A., Beebe, D.J., and Handelsman, J. (2025). Co-zorbs: Motile, multispecies biofilms aid transport of diverse bacterial species. Proceedings of the National Academy of Sciences 122, e2417327122. 10.1073/pnas.2417327122.

24. Stauffer, D., and Aharony, A. (1994). Introduction to percolation theory 2nd ed. (Taylor & Francis Group).

25. da Silva, C.R., Lyra, M.L., and Viswanathan, G.M. (2002). Largest and second largest cluster statistics at the percolation threshold of hypercubic lattices. Phys. Rev. E 66, 056107. 10.1103/PhysRevE.66.056107.

26. da Silva, C.R., Lyra, M.L., and Viswanathan, G.M. (2000). Boundary condition dependence of cluster size ratios in random percolation. Int. J. Mod. Phys. C 11, 1411– 1415. 10.1142/S0129183100001231.

27. Margolina, A., Herrmann, H.J., and Stauffer, D. (1982). Size of largest and second largest cluster in random percolation. Physics Letters A 93, 73–75. 10.1016/0375-9601(82)90219-5.

28. Trivedi, A., Gosai, J., Nakane, D., and Shrivastava, A. (2022). Design Principles of the Rotary Type 9 Secretion System. Front. Microbiol. 13, 845563. 10.3389/fmicb.2022.845563.

29. Zhulin, I.B., Nikolskaya, A.N., and Galperin, M.Y. (2003). Common extracellular sensory domains in transmembrane receptors for diverse signal transduction pathways in bacteria and archaea. J Bacteriol 185, 285–294. 10.1128/JB.185.1.285-294.2003.

30. Aravind, L., and Ponting, C.P. (1997). The GAF domain: an evolutionary link between diverse phototransducing proteins. Trends Biochem Sci 22, 458–459. 10.1016/s0968-0004(97)01148-1.

31. Pardoux, R., Dolla, A., and Aubert, C. (2021). Metal-containing PAS/GAF domains in bacterial sensors. Coordination Chemistry Reviews 442, 214000. 10.1016/j.ccr.2021.214000.

32. Spiro, S., and D’Autréaux, B. (2012). Non-Heme Iron Sensors of Reactive Oxygen and Nitrogen Species. Antioxidants & Redox Signaling 17, 1264–1276. 10.1089/ars.2012.4533.

33. Tucker, N.P., D’Autréaux, B., Yousafzai, F.K., Fairhurst, S.A., Spiro, S., and Dixon, R. (2008). Analysis of the Nitric Oxide-sensing Non-heme Iron Center in the NorR Regulatory Protein *. Journal of Biological Chemistry 283, 908–918. 10.1074/jbc.M705850200.

34. Tucker, N.P., D’Autréaux, B., Spiro, S., and Dixon, R. (2006). Mechanism of transcriptional regulation by the *Escherichia coli* nitric oxide sensor NorR. Biochem Soc Trans 34, 191–194. 10.1042/BST0340191.

35. Shrivastava, A., Rhodes, R.G., Pochiraju, S., Nakane, D., and McBride, M.J. (2012). Flavobacterium johnsoniae RemA Is a Mobile Cell Surface Lectin Involved in Gliding. J Bacteriol 194, 3678–3688. 10.1128/JB.00588-12.

36. Sankhe, G.D., Xing, J., Xiao, X., Buglino, J.A., Li, H., Zhulin, I.B., and Glickman, M.S. Ligand binding represses bacterial histidine kinase activity by inhibiting its dimerization. The FEBS Journal n/a. 10.1111/febs.70621.

37. Saito, K., Ito, E., Hosono, K., Nakamura, K., Imai, K., Iizuka, T., Shiro, Y., and Nakamura, H. (2003). The uncoupling of oxygen sensing, phosphorylation signalling and transcriptional activation in oxygen sensor FixL and FixJ mutants. Mol Microbiol 48, 373–383. 10.1046/j.1365-2958.2003.03446.x.

38. Monk, I.R., Shaikh, N., Begg, S.L., Gajdiss, M., Sharkey, L.K.R., Lee, J.Y.H., Pidot, S.J., Seemann, T., Kuiper, M., Winnen, B., et al. (2019). Zinc-binding to the cytoplasmic PAS domain regulates the essential WalK histidine kinase of Staphylococcus aureus. Nat Commun 10, 3067. 10.1038/s41467-019-10932-4.

39. Galperin, M.Y. (2006). Structural classification of bacterial response regulators: diversity of output domains and domain combinations. J Bacteriol 188, 4169–4182. 10.1128/JB.01887-05.

40. Baichoo, N., Wang, T., Ye, R., and Helmann, J.D. (2002). Global analysis of the Bacillus subtilis Fur regulon and the iron starvation stimulon. Mol Microbiol 45, 1613–1629. 10.1046/j.1365-2958.2002.03113.x.

41. Lim, C.K., Hassan, K.A., Tetu, S.G., Loper, J.E., and Paulsen, I.T. (2012). The Effect of Iron Limitation on the Transcriptome and Proteome of *Pseudomonas fluorescens* Pf-5. PLOS ONE 7, e39139. 10.1371/journal.pone.0039139.

42. Contreras-Moreno, F.J., Moraleda-Muñoz, A., Marcos-Torres, F.J., Cuéllar, V., Soto, M.J., Pérez, J., and Muñoz-Dorado, J. (2024). Siderophores and competition for iron govern myxobacterial predation dynamics. ISME J 18, wrae077. 10.1093/ismejo/wrae077.

43. Jo, J., Price-Whelan, A., and Dietrich, L.E.P. (2022). Gradients and consequences of heterogeneity in biofilms. Nat Rev Microbiol 20, 593–607. 10.1038/s41579-022-00692-2.

44. Werner, E., Roe, F., Bugnicourt, A., Franklin, M.J., Heydorn, A., Molin, S., Pitts, B., and Stewart, P.S. (2004). Stratified Growth in *Pseudomonas aeruginosa* Biofilms. Applied and Environmental Microbiology 70, 6188–6196. 10.1128/AEM.70.10.6188-6196.2004.

45. Yoon, S.S., Hennigan, R.F., Hilliard, G.M., Ochsner, U.A., Parvatiyar, K., Kamani, M.C., Allen, H.L., DeKievit, T.R., Gardner, P.R., Schwab, U., et al. (2002). *Pseudomonas aeruginosa* anaerobic respiration in biofilms: relationships to cystic fibrosis pathogenesis. Dev Cell 3, 593–603. 10.1016/s1534-5807(02)00295-2.

46. Sato, K., Naya, M., Hatano, Y., Kasahata, N., Kondo, Y., Sato, M., Takebe, K., Naito, M., and Sato, C. (2021). Biofilm Spreading by the Adhesin-Dependent Gliding Motility of Flavobacterium johnsoniae: 2. Role of Filamentous Extracellular Network and Cell-to-Cell Connections at the Biofilm Surface. Int J Mol Sci 22, 6911. 10.3390/ijms22136911.

47. Sato, K., Naya, M., Hatano, Y., Kondo, Y., Sato, M., Nagano, K., Chen, S., Naito, M., and Sato, C. (2021). Biofilm Spreading by the Adhesin-Dependent Gliding Motility of Flavobacterium johnsoniae. 1. Internal Structure of the Biofilm. International Journal of Molecular Sciences 22, 1894. 10.3390/ijms22041894.

48. Kearns, D.B. (2010). A field guide to bacterial swarming motility. Nat Rev Microbiol 8, 634– 644. 10.1038/nrmicro2405.

49. Moreau, A., Nguyen, D.T., Hinbest, A.J., Zamora, A., Weerasekera, R., Matej, K., Zhou, X., Sanchez, S., Rodriguez Brenes, I., Tai, J.-S.B., et al. (2025). Surface remodeling and inversion of cell-matrix interactions underlie community recognition and dispersal in Vibrio cholerae biofilms. Nat Commun 16, 327. 10.1038/s41467-024-55602-2.

50. Yoon, M.Y., Lee, K.-M., Park, Y., and Yoon, S.S. (2011). Contribution of Cell Elongation to the Biofilm Formation of *Pseudomonas aeruginosa* during Anaerobic Respiration. PLoS One 6, e16105. 10.1371/journal.pone.0016105.

51. Breskin, I., Soriano, J., Moses, E., and Tlusty, T. (2006). Percolation in Living Neural Networks. Phys. Rev. Lett. 97, 188102. 10.1103/PhysRevLett.97.188102.

52. Sahimi, M. (2023). Applications of Percolation Theory (Springer International Publishing) 10.1007/978-3-031-20386-2.

53. Silva, K.P.T., Yusufaly, T.I., Chellamuthu, P., and Boedicker, J.Q. (2019). Disruption of microbial communication yields a two-dimensional percolation transition. Phys. Rev. E 99, 042409. 10.1103/PhysRevE.99.042409.

54. Larkin, J.W., Zhai, X., Kikuchi, K., Redford, S.E., Prindle, A., Liu, J., Greenfield, S., Walczak, A.M., Garcia-Ojalvo, J., Mugler, A., et al. (2018). Signal Percolation within a Bacterial Community. cels 7, 137–145.e3. 10.1016/j.cels.2018.06.005.

55. Basaran, M., Yüce, T.C., Yaman, Y.I., Vetter, R., and Kocabas, A. (2022). Bacterial biofilms use chiral branches to escape crowded environments by tracking oxygen gradient. Preprint at arXiv, 10.48550/arXiv.2208.09730 https://doi.org/10.48550/arXiv.2208.09730.

56. Liu, J., Prindle, A., Humphries, J., Gabalda-Sagarra, M., Asally, M., Lee, D.D., Ly, S., Garcia-Ojalvo, J., and Süel, G.M. (2015). Metabolic codependence gives rise to collective oscillations within biofilms. Nature 523, 550–554. 10.1038/nature14660.

57. Asally, M., Kittisopikul, M., Rué, P., Du, Y., Hu, Z., Çağatay, T., Robinson, A.B., Lu, H., Garcia-Ojalvo, J., and Süel, G.M. (2012). Localized cell death focuses mechanical forces during 3D patterning in a biofilm. Proceedings of the National Academy of Sciences 109, 18891–18896. 10.1073/pnas.1212429109.

58. Claverie, C., Coppolino, F., Mazzuoli, M.-V., Guyonnet, C., Jacquemet, E., Legendre, R., Sismeiro, O., De Gaetano, G.V., Teti, G., Trieu-Cuot, P., et al. (2024). Constitutive activation of two-component systems reveals regulatory network interactions in Streptococcus agalactiae. Nat Commun 15, 9175. 10.1038/s41467-024-53439-3.

59. Magesh, S., Hurley, A.I., Nepper, J.F., Chevrette, M.G., Schrope, J.H., Li, C., Beebe, D.J., and Handelsman, J. (2024). Surface colonization by *Flavobacterium johnsoniae* promotes its survival in a model microbial community. mBio 15, e03428–23. 10.1128/mbio.03428-23.

60. Rhodes, R.G., Samarasam, M.N., Shrivastava, A., van Baaren, J.M., Pochiraju, S., Bollampalli, S., and McBride, M.J. (2010). *Flavobacterium johnsoniae* gldN and gldO Are Partially Redundant Genes Required for Gliding Motility and Surface Localization of SprB. Journal of Bacteriology 192, 1201–1211. 10.1128/JB.01495-09.

61. Sen, S., Vairagare, I., Gosai, J., and Shrivastava, A. (2025). RABiTPy: an open-source Python software for rapid, AI-powered bacterial tracking and analysis. BMC Bioinformatics 26, 127. 10.1186/s12859-025-06145-w.

62. Schindelin, J., Arganda-Carreras, I., Frise, E., Kaynig, V., Longair, M., Pietzsch, T., Preibisch, S., Rueden, C., Saalfeld, S., Schmid, B., et al. (2012). Fiji: an open-source platform for biological-image analysis. Nat Methods 9, 676–682. 10.1038/nmeth.2019.

63. Berg, S., Kutra, D., Kroeger, T., Straehle, C.N., Kausler, B.X., Haubold, C., Schiegg, M., Ales, J., Beier, T., Rudy, M., et al. (2019). ilastik: interactive machine learning for (bio)image analysis. Nat Methods 16, 1226–1232. 10.1038/s41592-019-0582-9.

64. Ershov, D., Phan, M.-S., Pylvänäinen, J.W., Rigaud, S.U., Le Blanc, L., Charles-Orszag, A., Conway, J.R.W., Laine, R.F., Roy, N.H., Bonazzi, D., et al. (2022). TrackMate 7: integrating state-of-the-art segmentation algorithms into tracking pipelines. Nat Methods 19, 829–832. 10.1038/s41592-022-01507-1.

65. Rhodes, R.G., Pucker, H.G., and McBride, M.J. (2011). Development and Use of a Gene Deletion Strategy for *Flavobacterium johnsoniae* To Identify the Redundant Gliding Motility Genes remF, remG, remH, and remI. Journal of Bacteriology 193, 2418–2428. 10.1128/jb.00117-11.

66. Eckroat, T.J., Greguske, C., and Hunnicutt, D.W. (2021). The Type 9 Secretion System Is Required for Flavobacterium johnsoniae Biofilm Formation. Frontiers in Microbiology 12.

67. Yang, Z., Zeng, X., Zhao, Y., and Chen, R. (2023). AlphaFold2 and its applications in the fields of biology and medicine. Sig Transduct Target Ther 8, 115. 10.1038/s41392-023-01381-z.

68. Blum, M., Andreeva, A., Florentino, L.C., Chuguransky, S.R., Grego, T., Hobbs, E., Pinto, B.L., Orr, A., Paysan-Lafosse, T., Ponamareva, I., et al. (2025). InterPro: the protein sequence classification resource in 2025. Nucleic Acids Res 53, D444–D456. 10.1093/nar/gkae1082.

69. Letunic, I., and Bork, P. (2026). SMART v10: three decades of the protein domain annotation resource. Nucleic Acids Res 54, D499–D503. 10.1093/nar/gkaf1023.

70. Meng, E.C., Goddard, T.D., Pettersen, E.F., Couch, G.S., Pearson, Z.J., Morris, J.H., and Ferrin, T.E. (2023). UCSF ChimeraX: Tools for structure building and analysis. Protein Science 32, e4792. 10.1002/pro.4792.

71. Kim, D., Paggi, J.M., Park, C., Bennett, C., and Salzberg, S.L. (2019). Graph-based genome alignment and genotyping with HISAT2 and HISAT-genotype. Nat Biotechnol 37, 907–915. 10.1038/s41587-019-0201-4.

72. Liao, Y., Smyth, G.K., and Shi, W. (2014). featureCounts: an efficient general purpose program for assigning sequence reads to genomic features. Bioinformatics 30, 923–930. 10.1093/bioinformatics/btt656.

73. Robinson, M.D., McCarthy, D.J., and Smyth, G.K. (2010). edgeR: a Bioconductor package for differential expression analysis of digital gene expression data. Bioinformatics 26, 139–140. 10.1093/bioinformatics/btp616.

74. Virtanen, P., Gommers, R., Oliphant, T.E., Haberland, M., Reddy, T., Cournapeau, D., Burovski, E., Peterson, P., Weckesser, W., Bright, J., et al. (2020). SciPy 1.0: fundamental algorithms for scientific computing in Python. Nat Methods 17, 261–272. 10.1038/s41592-019-0686-2.

75. McKinney, W. (2010). Data Structures for Statistical Computing in Python. In, pp. 56–61. 10.25080/Majora-92bf1922-00a.

76. PANNZER—A practical tool for protein function prediction - PMC https://pmc.ncbi.nlm.nih.gov/articles/PMC8740830/.

77. Chiu, C.-L., Clack, N., and the napari community (2022). napari: a Python Multi-Dimensional Image Viewer Platform for the Research Community. Microscopy and Microanalysis 28, 1576–1577. 10.1017/S1431927622006328.

78. Hartmann, R., Jeckel, H., Jelli, E., Singh, P.K., Vaidya, S., Bayer, M., Rode, D.K.H., Vidakovic, L., Díaz-Pascual, F., Fong, J.C.N., et al. (2021). Quantitative image analysis of microbial communities with BiofilmQ. Nat Microbiol 6, 151–156. 10.1038/s41564-020-00817-4.

79. Ahrens, J., Geveci, B., and Law, C. (2005). ParaView: An End-User Tool for Large-Data Visualization. In Visualization Handbook (Elsevier), pp. 717–731. 10.1016/B978-012387582-2/50038-1.

